# The ecology of sexual conflict: behaviorally plastic responses to temperature variation in the social environment can drastically modulate male harm to females

**DOI:** 10.1101/429514

**Authors:** Roberto García-Roa, Valeria Chirinos, Carazo Pau

**Author notes:** Author for correspondence **Contact Information:** Roberto García-Roa, Behaviour and Evolution group, Ethology lab, Cavanilles Institute of Biodiversity and Evolutionary Biology, University of Valencia, Valencia, Spain; Valeria Chirinos, Behaviour and Evolution group, Ethology lab, Cavanilles Institute of Biodiversity and Evolutionary Biology, University of Valencia, Valencia, Spain; Pau Carazo, Behaviour and Evolution group, Ethology lab, Cavanilles Institute of Biodiversity and Evolutionary Biology, University of Valencia, Valencia, Spain. **Authorship:** RGR and PC conceived this study. RGR and PC designed the experiments, and RGR & VC-B conducted them with punctual help from PC. RGR and PC analysed arising results and wrote the manuscript. **Data accessibility statement:** once the manuscript is accepted, the data supporting the results will be archived in an appropriate public repository such as Dryad or Figshare and the data DOI will be included at the end of the article.

## Abstract

Sexual conflict is a fundamental driver of male/female adaptations, an engine of biodiversity, and a crucial determinant of population viability. For example, sexual conflict frequently leads to behavioural adaptations that allow males to displace their rivals, but in doing so harm those same females they are competing to access. Sexual conflict via male harm hence not only deviates females from their fitness optimum, but can decrease population viability and facilitate extinction. Despite this prominent role, we are far from understanding what factors modulate the intensity of sexual conflict, and particularly the role of ecology in mediating underlying behavioural adaptations. In this study we show that, in *Drosophila melanogaster*, variations in environmental temperature of ±4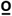C (within the natural range in the wild) decrease male harm impact on female fitness by between 45–73%. Rate-sensitive fitness estimates indicate that such modulation results in an average rescue of population productivity of 7% at colder temperatures and 23% at hotter temperatures. Our results: a) show that the thermal ecology of social interactions can drastically modulate male harm via behaviourally plasticity, b) identify a potentially crucial ecological factor to understand how sexual conflict operates in nature, and c) suggest that behaviourally plastic responses can lessen the negative effect of sexual conflict on population viability in the face of rapid environmental temperature changes.

## Introduction

The classic perception of sexual reproduction as a cooperative endeavour has been shattered over the last decades. There is now ample evidence to show that the fitness of one sex frequently raises at the expense of the other, so that male and female evolutionary interests cannot be simultaneously maximized (Parker 1979; Arnqvist & Rowe 2013; Parker 2014). This process, termed ‘sexual conflict’, is particularly important in polygynous species but applies to most (if not all) mating systems (Hosken *et al*. 2008), and is hence rampant across the tree of life (Parker & Partridge 1998; Stutt & Siva-Jothy 2001; Parker 2006; Bonduriansky & Chenoweth 2009; Kalbitzer *et al*. 2017). Sexual conflict can reduce male and female fitness via gender load (Arnqvist & Tuda 2009; Berger *et al*. 2016), and can additionally curtail female fitness when strong sexual selection in males leads to adaptations that harm females (i.e. male harm), either as a direct or collateral consequence of male-male competition (Hall *et al*. 2008; Michalczyk *et al*. 2011; Tobias *et al*. 2012). Male harm is well documented and widespread in nature, whereby males increase their own reproductive output at the expense of females by harassing, punishing or coercing them during reproduction (Clutton-Brock & Parker 1995), for example due to traumatic inseminations (Crudgington & Siva-Jothy 2000) or the transfer of toxic proteins in their ejaculates (Chapman 2001; Wigby & Chapman 2005). Crucially, via gender load and/or male harm, sexual conflict may reduce population productivity and profoundly impact population viability (Le Galliard *et al*. 2005; Rankin *et al*. 2007; Berger *et al*. 2016). In contrast, sexual conflict may also act as an engine of speciation (Maan & Seehausen 2011; Gavrilets 2014), as it can contribute to genetic divergence between populations and the evolution of reproductive isolation ((Parker & Partridge 1998; Gavrilets 2000); reviewed in (Gavrilets 2014)). Understanding what factors modulate the intensity of sexual conflict is hence a priority (Dean *et al*. 2007; Pizzari *et al*. 2007; Cox & Calsbeek 2010; Eldakar *et al*. 2010; Pizzari *et al*. 2015).

Recent work has emphasized that sexual conflict must be understood in its ecological setting (Martinossi-Allibert *et al*. 2017; Perry *et al*. 2017b; De Lisle *et al*. 2018; Gomez-Llano *et al*. 2018; Perry & Rowe 2018). On the one hand, several studies over the last few years have shown that intralocus sexual conflict can be strongly modulated by the environment, so that inter-sexual correlations in fitness can change significantly across environments (Long *et al*. 2012; Berger *et al*. 2014; Punzalan *et al*. 2014); but see (Delcourt *et al*. 2009; Punzalan *et al*. 2014; Martinossi-Allibert *et al*. 2018a). On the other, a few studies have also shown that inter-locus sexual conflict can be similarly affected by the environment (Martinossi-Allibert *et al*. 2018b). For example, using *D. melanogaster*, Arbuthnott *et al*. (2014) showed that evolution of male harm and female resistance is conditioned by the environment, and Yun *et al*. showed that the physical environment (i.e. small and simple vs. large and complex) drastically affects sexual interactions and male harm in the same species (Yun *et al*. 2017). Generally speaking, sexually antagonistic selection is predicted to increase as a population is better adapted to its environment and in stable environments (and vice versa), a prediction that was supported by a meta-analysis of over 700 studies measuring sex-specific phenotypic selection (De Lisle *et al*. 2018).

Temperature is probably one of the best known ecological factors influencing the biology of animal systems (Kristensen *et al*. 2008; Olsson *et al*. 2011; Monteiro *et al*. 2017; Nowakowski *et al*. 2018; Parrett & Knell 2018). Especially relevant for ectotherms, the thermal environment can impact a wealth of sexually selected morphological traits and behaviours (Stillwell & Fox 2007; Punzalan *et al*. 2008; Llusia *et al*. 2013; Monteiro *et al*. 2017), as well as the sex-specific costs and benefits of reproduction (Grazer & Martin 2012), operational sex ratios (Kvarnemo 1996), potential reproductive rates (Kvarnemo 1994; Ahnesjo 1995) and/or spatio-temporal distributions (Møller 2004), which ultimately are all susceptible to modulate both the nature and intensity of sexual conflict (Wigby & Chapman 2005; Eldakar *et al*. 2009; Svensson *et al*. 2009; Cox & Calsbeek 2010). Indeed, a recent study showed that intralocus sexual conflict decreases at stressful temperatures in one of two natural populations of seed beetles (*Callosobruchus maculatus*), purportedly due to the increased importance of mutations with condition dependent fitness effects (Berger *et al*. 2014). In other words, alleles with sexually antagonistic effects in a benign environment (i.e. positive for males and negative for females) tend to have negative effects for both sexes in a stressful environment (Berger *et al*. 2014). This study nicely demonstrate how temperature can modulate intra-locus sexual conflict, reducing gender load in populations subject to a stressful environment precisely when adaptation via sexual selection is most needed. Importantly, beyond its stressful effects in extreme thermal environments, temperature can be expected to impact sexual selection processes, including sexual conflict via male harm, within the normal range of temperatures experienced by organisms in their environment. In nature, adult, sexually mature organisms are bound to experience daily, intra-seasonal and inter-seasonal fluctuations in temperature that can affect a suite of physiological and behavioural reproductive parameters (e.g. metabolic rate, spermatogenesis, operational sex ratios, potential reproductive rates, etc.) that could in turn modulate sexual conflict. In accordance with this idea, Perry *et al*. (2017a) found temperature to be among the ecological factors modulating inter-population variation in a sexually antagonistic evolutionary arms race (in the water strider *G. incognitus*). Similarly, De Lisle et al. (De Lisle *et al*. 2018) found that the strength of sexually antagonistic section is partly explained by microclimatic conditions such as temperature and altitude.

Our aim in this paper was to explore whether environmental temperature shifts during reproductive interactions, within a range frequently experienced by natural populations in the wild, can modulate sexual conflict intensity through male harm. Specifically, in this paper we asked whether such modulation may happen via behaviourally plastic changes. Studying how behavioural plasticity may modulate sexual conflict in response to ecological fluctuations is critical. First, to understand sexual conflict levels and their impact on male/female fitness and population viability in natural conditions, where such fluctuations are common. Second, because behavioural plasticity is the first line of defence against a changing environment, and hence key to understand a population’s potential for evolutionary rescue. For this purpose, we used *D. melanogaster*, a model system in sexual conflict studies with high levels of male-male competition and sexual conflict driven by male harm (Chapman *et al*. 1995; Wigby & Chapman 2004), and where female reproductive behaviours and sexual conflict mechanisms have been very well studied (Rice 1996; Wigby & Chapman 2005; Bretman *et al*. 2009; Manier *et al*. 2010).

In a first experiment, we reared flies under standard conditions (i.e. 25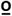C) and then exposed them (as sexually mature adults) to different temperatures within a range experienced by *Drosophila melanogaster* populations in the wild (i.e. 21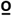C, 25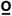C and 29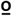C); and at which they are reproductively active both in the lab and in the field (Hoffmann *et al*. 2002; Ashburner *et al*. 2005). To gauge male harm levels, at each temperature we examined the decrease in focal female fitness (i.e. lifespan, lifetime reproductive success and reproductive senescence) in a high sexual conflict context (i.e. three males competing over access to one female) vs. a low sexual conflict context (i.e. one male and one female). We found that the strongest decrease in female fitness (i.e. male harm levels) occurred at 25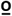C (the temperature at which experimental flies were reared and the average temperature at which this population is maintained in the lab). Modulation of male harm at 21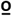C and 29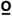C was so high that the strong fitness advantage of female flies at 25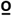C in a low sexual conflict context (i.e. >25% increase in lifetime reproductive success at 25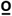C *vs*. 21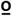C/29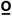C) completely disappeared in a high sexual conflict context. We replicated this experiment to focus on changes in male (i.e. sexual harassment levels and re-mating rates) and female behaviours (i.e. rejection rates), with the aim of examining potential behavioural mechanisms underlying these effects. We observed that mating frequency and female rejections increased relatively more in the high sexual conflict context at 25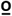C, which reinforces our previous finding that male harm is greater at this temperature. Our results demonstrate that behaviourally plastic responses to normal shifts in the environmental temperature of social interactions drastically modulate male harm levels. We discuss the consequences that this finding has for our understanding of sexual conflict levels in the wild, as well as the possible implications for population viability in the face of rapid environmental changes.

## Material and methods

### Stock culture and maintenance

All experiments were conducted on individuals from a laboratory-adapted wild-type stock of *D. melanogaster* maintained in an outbred population since 1970 (Partridge & Farquhar 1983). This population is maintained in the laboratory with overlapping generations at 25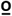C, at 50–60% humidity and a 12:12 h light:dark cycle. Experimental flies were collected following the same procedure. Yeasted grape-juice agar plates were introduced into stock cultures to induce female oviposition. Eggs were then collected and placed in bottles with standard food to be incubated at 25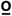C at a controlled density (Clancy & Kennington 2001). Emerging virgin flies were sexed and isolated at 25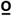C during 2–3 days. After this, they were individually allocated to the different temperature treatments 48 hours prior to the beginning of the experiment.

### Experiment 1: Temperature effects on levels of male harm

To investigate whether sexual conflict is affected by temperature we set-up a full factorial design in which we measured the lifespan and fitness of female flies under low sexual conflict (i.e. one male and one female in a single vial) *vs*. high sexual conflict (i.e. 3 males competing over access to one female in a single vial), across three thermal treatments: 21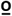C, 25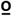C and 29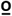C. We also included a set of isolated females (one female in a single vial) at 21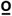C, 25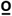C and 29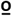C as a control.

Once the experiment started, we transferred focal flies to fresh vials twice a week using gentle CO_2_ anaesthesia (i.e. a short, ∼2s puff of CO_2_ from which flies recovered within the minute). Vials containing female eggs were incubated at 25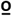C for 16 days and then frozen at –21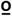C until offspring could be counted. At the end of the first week, we isolated focal males to estimate treatment effects on male lifespan, and introduced new males of the same age to continue with the experiment. After the second week, competitor males were discarded and replaced by young 2–4d-old virgin males every 10–12 days (at the same time for all treatments). We kept a stock of flies during each round of collection to replace dead competitor flies. Flies were kept under these conditions until all focal females and males died. We started the experiment with 450 females (50 per each temperature*sexual conflict level treatment) and 600 males (150 per each temperature*high sexual conflict level and 50 per each temperature*low sexual conflict level). Final sample sizes were: a) at 21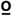C: 47 females and 132 males for high sexual conflict, 50 females and 49 males for low sexual conflict, and 50 females for isolation (i.e. control) treatments; b) at 25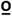C: 50 females and 129 males for high sexual conflict, 50 females and 44 males for low sexual conflict, and 49 females for isolation treatments; c) at 29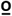C: 49 females and 126 males for high sexual conflict, 48 females and 46 males for low sexual conflict, and 49 females for isolation treatments. Differences between starting and final sample sizes across treatments are due to escaped flies during the experiment, and/or flies that died before reproducing (but note these were taken into account –i.e. right censored– in survival analyses).

### Experiment 2: Behavioural assays

To examine behavioural mechanisms that might underlie the fitness effects observed in our first experiment, we conducted a series of behavioural observation trials in which we compared the reproductive behaviour of male and female flies subject to the same factorial design imposed in experiment 1. All focal flies were collected as in experiment 1. At 2–3 days post-eclosion, focal flies were introduced into their randomly assigned temperature treatment, 12h immediately prior to the start of mating trials (i.e. at lights-off the day prior to behavioural trials). Behavioural observation had to be conducted in the same temperature control room, so trials at 21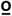C, 25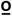C and 29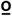C had to be conducted in three consecutive days, in randomized order (i.e. 25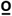C, 21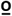C and 29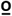C). Sample sizes were of 65 vials per treatment combination (i.e. 130 vials per day/temperature treatment, for a total of 390 vials and 1170 flies). We measured the following behaviours: a) courtship intensity (number of courtships experienced by a female per minute), b) latency to the first mating, c) mating duration (first mating), d) re-mating rates (i.e. number of total matings during the observation period) and e) female rejection behaviours (Bastock & Manning 1955; Connolly & Cook 1973). Observations started at lights-on (i.e. 10am), and lasted for 8h, during which time reproductive behaviours were continuously recorded using scan sampling of vials (one ∼12 minutes scan every 15 minutes). Scans consisted in observing all vials in succession for 3s each, and recording all occurrences of the behaviours listed above that were observed within such time frame. We interspersed these ‘behavioural’ scans with very quick (< 1 min) ‘copula’ scans where we rapidly swept all vials for copulas at the beginning, in the middle and at the end of each ‘behavioural’ scan. This strategy ensured that we recorded all matings that took place during our 8h observation trials because each vial was scanned for copulas at least once every <10 minutes, while successful matings typically take 15 minutes or more in our population of *D. melanogaster*.

### Statistical analyses

We examined the effect of temperature (i.e. 21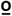C, 25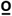C and 29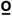C), sexual conflict level (i.e. high vs. low) and their interaction on the reproductive behaviours measured using GLMs with temperature, sexual conflict level and their interaction as fixed factors. The variables courtship rate (i.e. courtships per minute), mating time (i.e. time in copula per minute) and mating duration (i.e. duration of the first mating in the vial) were winsorized at α = 0.05 to control for the presence of outliers, and then tested using GLMs with Gaussian error distributions. Latency to mate exhibited a right skew, and was square root transformed prior to running a GLM with a Gaussian error distribution. Female rejection behaviours were analysed in two different ways. First, we examined rejection rate (i.e. female rejections per minute). We used a Gamma distribution for this analysis as this variable was continuous and zero-inflated. Second, we also used a binomial GLM to examine temperature, sexual conflict level and temperature*sexual conflict level effects on the proportion of courtships that were rejected by females. Finally, we analysed the total number of matings recorded across the 8h observation period (i.e. re-mating rate) using Poisson and Quasipoisson GLM models for count data. As a complementary analysis, we also examined the variable mating rate (i.e. number of matings recorded over the 8h observation period divided by the exact observation period in each vial). Across behavioural observation assays, each of the three days we run tests it took us 15–20 minutes to poot in all the experimental flies into the experimental vials, which means there was a maximum difference of ∼20 minutes in the overall time flies had to mate across vials. Given treatments were perfectly counterbalanced across vials, such minor differences in overall observation time (i.e. < 5% of total observation time) are not expected to bias our results, but we nevertheless decided to examine the variable mating rate too. Due to a strong left skew, we used a boxcox transformation (Quinn & Keough 2002) prior to running a GLM with a Gaussian error distribution. In all of the cases above we checked data for heteroscedasticity and normality assumptions prior to fitting models, and all fitted models were subsequently validated. Reported p-values are one-way. All analyses were conducted in R.3.2.4 (R Core Team. 2014. R: A language and environment for statistical computing. R Foundation for Statistical Computing 2014).

## Results

### Survival, reproductive senescence and lifetime reproductive success

We detected a significant temperature*sexual conflict level interaction for female lifespan (F _8,431_=26.64, *P* < 0.001; Fig. 1a), as well as main temperature (F _2,286_=190.59, *P* < 0.001) and sexual conflict level effects (F _2,286_=381.99, *P* < 0.001). The interaction dropped to marginally non-significant when we removed isolated females from statistical comparisons (F_5,286_=2.87, *P* = 0.058), but main effects remained significant (temperature: F_2,286_=101.89, *P* < 0.001; sexual conflict level: F_1,286_=173.68, *P* < 0.001). We did not detect a significant temperature*sexual conflict level interaction for male lifespan (F_5,252_=1.86, *P* = 0.158) but both main effects were significant (temperature: F_2,252_=205.76, *P* < 0.001; sexual conflict level: F_1,252_=32.47, *P* < 0.001). Survival analysis by means of a Cox proportional hazards model did detect significant temperature*sexual conflict level interactions for both females (d.f. = 8, z = 725.9, *P* < 0.001; Fig. S1) and males (d.f. = 5, z = 391.1, *P* < 0.001).

**Figure 1.**
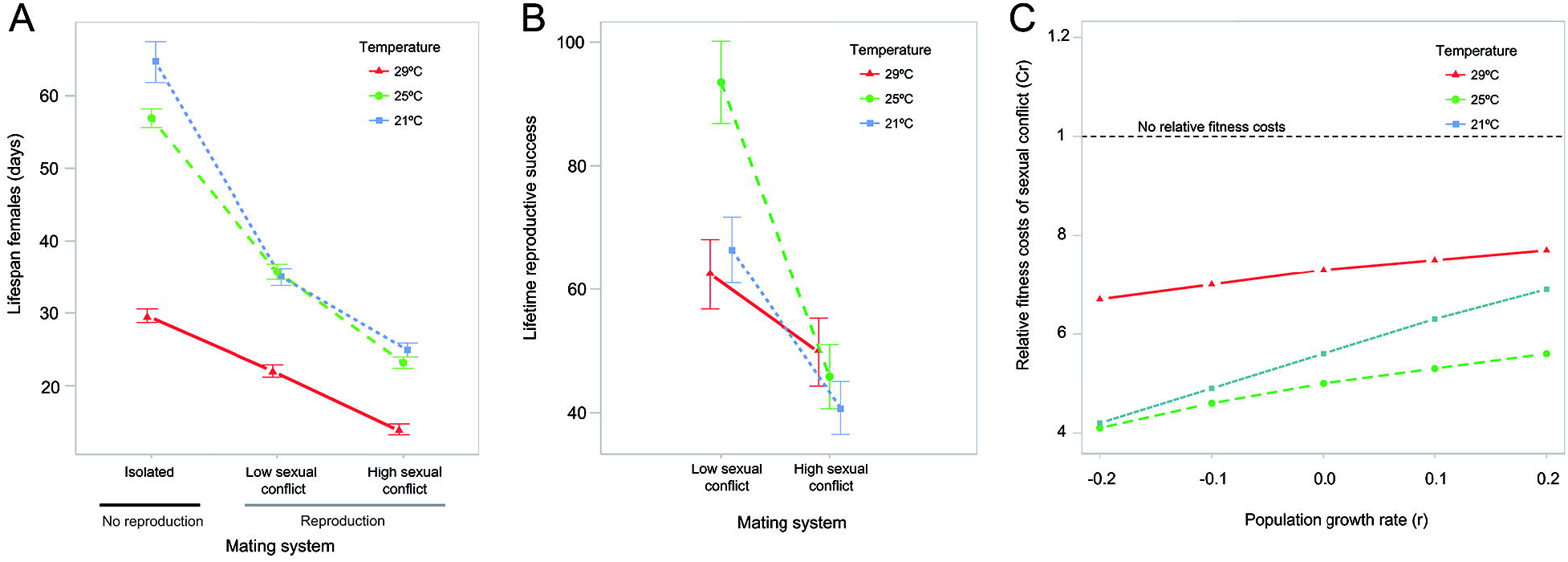
Fitness effects of sexual conflict across temperature and sexual conflict level treatments (experiment 1): a) decrease in female lifespan (i.e. actuarial senescence) with high sexual conflict, low sexual conflict, and in isolation, b) decrease in lifetime reproductive success (i.e. total offspring produced) at high vs. low sexual conflict, and c) relative population fitness costs of high (vs. low) sexual conflict for different population growth rates (note the dashed horizontal line marks the isoline for no fitness costs at high vs. low sexual conflict). Data provided in mean ± sem.

In addition, the interaction between temperature and sexual conflict level was significant for reproductive ageing (Chi = 13.493, df = 2, *P* = 0.001). To check whether this interaction was driven by reproductive senescence at 21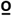C, as suggested by the interaction plot (Fig. S2), we re-fitted statistical models separately for the three different temperature treatments. We found a significant effect of reproductive senescence (i.e. time*sexual conflict level) at 21 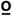C (GLMM; Chi = 4.24, df = 1, *P* = 0.039), but not at 25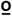C (GLMM; Chi = 1.69, df = 1, *P* = 0.193), or 29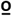C (GLMM; Chi =3.56, df = 1, *P* = 0.058; marginally non-significant).

Finally, we found a significant temperature*sexual conflict level interaction for female lifetime reproductive success (F_5,294_= 5.23, *P* = 0.005; Fig. 1b), as well as for the main effects of temperature (F_2,294_= 4.98, *P* = 0.007) and sexual conflict level (F_1,294_= 41.02, *P* < 0.001). To check whether, as suggested by the interaction plot (Fig. 1b), this interaction was driven by a stronger decrease (in the high vs. low sexual conflict treatment) in LRS at 25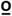C, we run models separately for each temperature level. Sexual conflict level affected the lifetime reproductive success at 21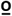C (F_1,98_= 13.98, *P* < 0.001, estimate = –25.6 ± 6.84) and at 25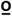C (F_1,98_= 32.03, *P* < 0.001, estimate = –47.68 ± 8.4), but not at 29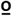C (F_1,98_= 2.57, *P* = 0.112, estimate = –12.64 ± 7.88). Hence, the decrease in LRS with sexual conflict was between 2 and 4 times larger at 25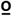C than at 21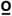C and 29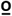C. We found qualitatively identical results when examining rate-sensitive fitness estimates (w_ind_, r = 0, (Edward *et al*. 2011); Fig 1c) across different population growth rates (i.e. r = –0.2 to 0.2; see Supp. Mat.).

### Behavioural assays

We detected significant temperature*sexual conflict level effects for courtship rate (F_2,384_ = 312.17, *P* < 0.001), as well as main temperature (F_2,386_ = 79.41, *P* < 0.001) and sexual conflict level effects (F_1,388_ = 104.38, *P* < 0.001; see Fig. 2a). Similarly, we detected significant temperature*sexual conflict level effects for latency to mate (F_2,361_ = 9.81, *P* < 0.001), as well as main temperature (F_2,363_ = 222.11, *P* < 0.001) and sexual conflict level effects (F_1,365_ = 36.71, *P* < 0.001; see Fig. 2b). In contrast, for mating duration we did not detect a significant temperature*sexual conflict level interaction (F_2,357_ = 2.443, *P* = 0.088), but we did detect significant temperature (F_2,359_ = 89.528, *P* < 0.001) and sexual conflict level effects (F_1,361_ = 7.235, *P* = 0.008; see Fig. 2c). As for female refection behaviour, we also detected a clear significant temperature*sexual conflict interaction for female rejection rate (F2,384 = 8.305, *P* = 0.004), and main effects for both temperature (F2,386 = 5.50, *P* = 0.004) and sexual conflict level (F2,388 = 16.80, *P* < 0.001; see Fig. 3a). Regarding re-mating behaviour, we detected a highly significant temperature*sexual conflict level interaction (F_2,384_ = 5.081, *P* = 0.007), as well as main temperature (F_2,386_ = 57.46, *P* < 0.001) and sexual conflict level effects (F_1,388_ = 59.46, *P* < 0.001; see Fig. 3b). Analysing mating rate instead yielded very similar results: temperature*sexual conflict level interaction (F_2,358_ = 5.081, *P* = 0.007), temperature (F_2,360_ = 57.46, *P* < 0.001) and sexual conflict level (F_1,362_ = 59.46, *P* < 0.001).

**Figure 3.**
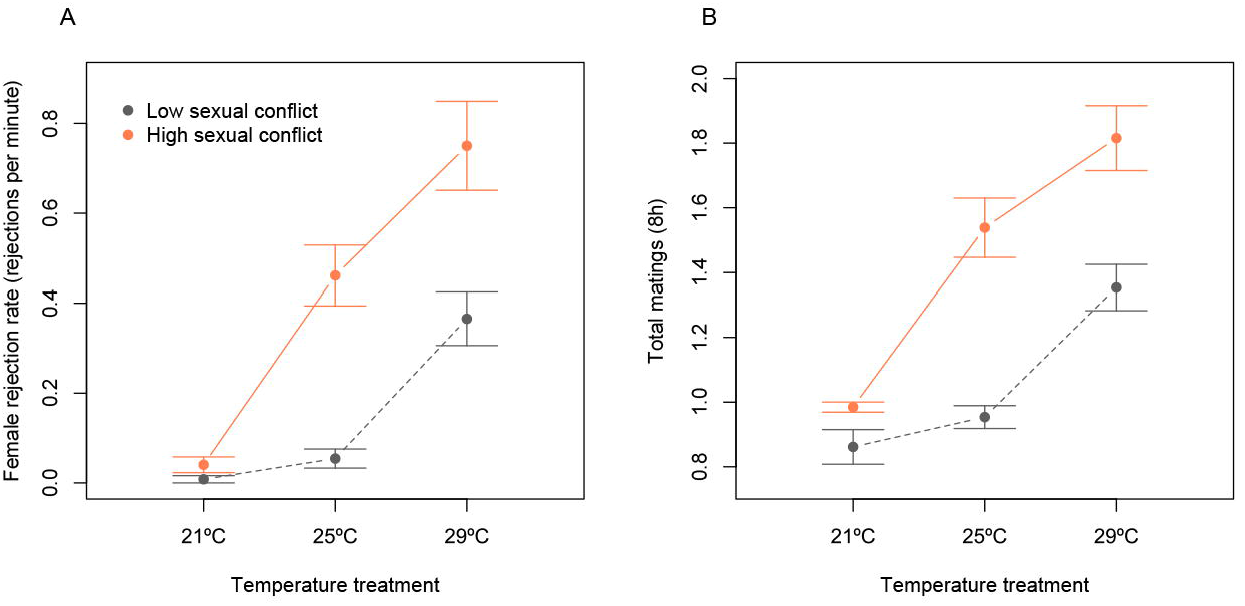
Behaviourally plastic differences in reproductive behaviour across temperature and sexual conflict level treatments (experiment 2): a) female rejection rate (i.e. rejections per minute) and b) total number of matings across the 8h of observations (i.e. re-mating rate). Data provided in mean ± sem.

**Figure 2.**
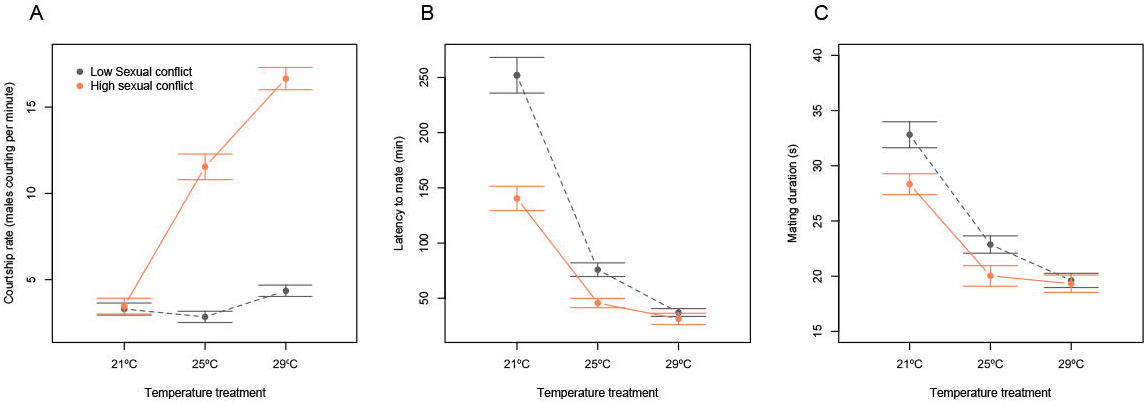
Behaviourally plastic differences in reproductive behaviour across temperature and sexual conflict level treatments (experiment 2): a) courtship rate (i.e. courtships per minute), b) latency to mate (in minutes), c) mating duration (in minutes). Data for (a) and (b) is provided in mean ± sem.

## Discussion

In this study, we show that variation in the environmental temperature experienced by adults during social interactions, within a range of variation where they are reproductively active and that is typically experienced by flies in the field (Hoffmann *et al*. 2002; Ashburner *et al*. 2005), can drastically modulate male harm. Results from behavioural assays further suggest that these effects cannot be explained by changes in male sexual harassment to females, but rather by temperature modulation of re-mating frequency and/or mating costs. Altogether, we provide evidence that temperature can modulate sexual conflict via male harm, most likely via behaviourally plastic non-linear effects on different underlying proximate mechanisms. These findings show that considering natural variation in the social thermal environment can be critical to understand sexual conflict in wild populations, and its consequences in terms of female productivity and population viability in the face of rapid environmental changes. The latter result fits well with recent studies (Berger *et al*. 2014; Martinossi-Allibert *et al*. 2018b) to suggest that the negative effect of sexual conflict on population viability can be significantly reduced in response to episodes of rapid environmental change (e.g. global warming), due to behaviourally plastic changes that seem to rescue population productivity.

### Effect of temperature on male harm (and its consequences for population viability)

For decades, investigations have ignored the potentially critical feedback that environmental temperature can have on sexual conflict. More recently, Perry *et al*. (2017) and De Lisle et al. (De Lisle *et al*. 2018) found temperature to be one of several ecological factors influencing a sexually antagonistic evolution in wild populations, and Berger *et al*. (2014) showed that stress induced by an extreme temperature treatment reduced intralocus sexual conflict (and thus gender load) in *C. maculatus* seed beetles. In this study, we evaluated whether temperature shifts similar to those experienced by flies in the field modulate the degree of male harm, and found that temperature does indeed drastically modulate male harm levels. In a low sexual conflict scenario, females had considerably higher fitness (> 25% advantage in LRS; Fig. 1) at 25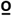C, their optimal average temperature, than at 21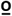C and 29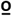C. However, this advantage all but disappeared in a high sexual conflict scenario, due to a decrease in female fitness via male harm that was roughly 2 and 4 times higher at 25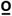C than at 21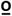C and 29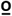C (Fig. 4). Rate-sensitive estimates, calculated across a wide range of population growth scenarios, indicate that such modulation of male harm results in an average rescue of population productivity under high sexual conflict of 7% (at colder temperatures) and 23% (at hotter temperatures; Fig. 1). At all three temperature treatments, male harm imposed greater costs in decreasing populations, which reflects that part of the fitness costs to females are differentially paid later in life (i.e. via the combined effect of actuarial and reproductive senescence; Fig. 1). Curiously, late-life fitness effects were more evident at 21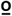C, where we found clear evidence of reproductive senescence. The fact that male harm significantly increased reproductive senescence at low (but not high) temperatures is extremely interesting, as it suggests temperature does not have linear effects on underlying male harm mechanisms (more on this below). Similarly, our results suggest that reproduction costs in terms of actuarial ageing (i.e. how much reproduction decreases female lifespan) are more marked at 21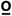C and 25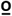C than at 29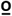C (Fig. 1), which again points towards non-linear effects of temperature on ageing processes.

**Figure 4.**
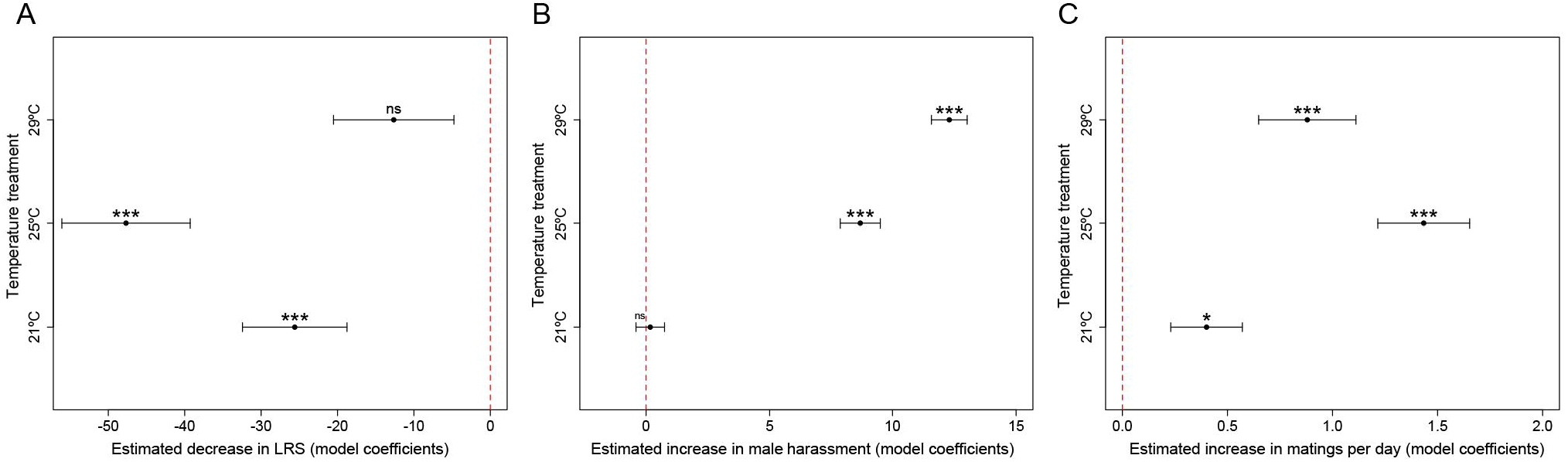
Model coefficient estimates (± sem) for the estimated variation in LRS (a), male harassment to females (b) and in the number of matings per day (c) that ensue with high (vs. low) sexual conflict, across the different temperature treatments. Estimates are coefficients from GLMs fitted separately for the three temperature treatments. LRS and male harassment estimates are for raw data (i.e. estimated decrease in the number of offspring –LRS– and courtship rate –courtship events per minute–), while estimates for number of matings are on a log link scale (i.e. quasipoisson GLM on the number of matings per 8h observation period).

The above results are relevant on three fronts. First, they constitute the first solid evidence, to our knowledge, that male harm can be modulated by temperature. This finding fits with recent studies in identifying ecology as an important factor for sexual conflict (e.g. Arbuthnott *et al*. 2014; Perry *et al*. 2017; see also Yun *et al*. 2017). In particular, it underscores the potentially crucial importance that temperature may play in modulating sexual conflict across taxa (De Lisle *et al*. 2018), and particularly so in ectotherms (Arbuthnott *et al*. 2014; Berger *et al*. 2014; Perry *et al*. 2017b). We now have direct evidence that temperature can modulate sexual conflict by decreasing both gender load (Berger *et al*. 2014) and male harm levels (this study). In addition, in a very recent study, Martinossi-Allibert et al. (Martinossi-Allibert *et al*. 2018b) evolved seed beetles under three alternative mating regimes (i.e. monogamy, polygamy or male-limited selection), and then tested in their ancestral vs. a stressful environment (i.e. at an elevated temperature or a new host). Interestingly, they found a trend whereby the general costs of socio-sexual interactions (i.e. including inter- and intra-sexual interactions) tended to be lower at elevated temperatures in both monogamy and polygamy (albeit not in male-limited evolution lines), which is consistent with a reduction in inter-locus sexual conflict (Martinossi-Allibert *et al*. 2018b).

Second, in this study we found drastic modulation of male harm in response to relatively mild fluctuations in the temperature (± 4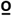C) experienced by adult flies during social interactions, while the only other precedents (Berger *et al*. 2014; Martinossi-Allibert *et al*. 2018b) focused on the effect of an extreme rise in temperature (i.e. stressful environment) across the whole developmental period (i.e. from egg to adult). Variations of the magnitude imposed in our study and higher are typically experienced by wild *D. melanogaster* adult flies both inter-seasonally, intra-seasonally and, more importantly, within a single day, and fall well within the temperature range at which flies are reproductively active in the field and in the lab (Hoffmann *et al*. 2002; Ashburner *et al*. 2005; Chen *et al*. 2012). Hence, our results suggest that acknowledging the effects of temperature might be crucial to understand the consequences of male harm in nature, as well as differences in male harm levels across time, populations and taxa. They also add to recent studies to suggest that we need to bring a sharper focus into the question of how natural fluctuations in the ecology of the social reproductive context may affect sexual conflict and sexual selection processes at large (see Yun *et al*. 2017).

Third, along with Berger *et al*. (2014) and Martinossi-Allibert et al. (Martinossi-Allibert *et al*. 2018b), our finding that behaviourally plastic responses to temperature shifts can modulate sexual conflict, and more specifically the population costs derived from sexual conflict, may have direct implications for our understanding of how rapid temperature change impacts population viability. In conjunction, these studies suggest that when a population experiences a shift away from their optimal environmental temperature, sexual conflict is relaxed via behaviourally plastic changes that result both in reduced gender load and male harm levels. This is in accordance with sexually antagonistic theory (De Lisle *et al*. 2018) and, in practice, means populations facing a thermal shift would be spared some of the sexual conflict costs in terms of both productivity (i.e. population growth rate) and evolvability (i.e. gender load), which ultimately translates into a higher probability of evolutionary rescue. It is important to note that studies have so far been conducted in insects, which may be particularly permeable to these effects due to the high sexual conflict levels generally reported in insects and to the fact that they are small ectotherms (and hence are more affected by temperature). A priority for future studies should thus be to expand these findings to other taxa, including endotherms.

### What mechanisms underlie temperature modulation of male harm in Drosophila melanogaster?

We replicated our experiment to examine the type of behavioural mechanisms that might underlie temperature modulation of male harm. In *D. melanogaster*, male harm occurs mainly via sexual harassment (Long *et al*. 2009) and/or toxic components in the male ejaculate (Chapman *et al*. 1995; Wigby & Chapman 2004, 2005). We found that temperature drastically increased male harassment of females in general, and that sexual harassment increased relatively more in the high sexual conflict scenario at 29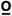C than at 25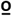C, and at 25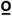C than at 21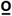C (see Fig. 2 and 4). Given that male harm increased more at 25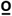C than at lower and higher temperatures, and that if anything we found male harm to be higher at 21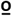C than at 29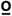C, sexual harassment fails to explain our results. In contrast, at 25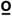C we detected a relatively higher increase in mating rates than at lower or higher temperatures (Fig. 3), which suggests increased sexual conflict costs are at least partly associated with higher re-mating rates and/or mating costs.

A growing body of literature has examined the important role that ejaculates play on reproductive behaviour and physiology of males and females, as well as on their fitness (reviewed in (Birkhead & Møller 1998; Simmons 2005; Simmons & Fitzpatrick 2012; Smith 2012)). In *D. melanogaster*, many of the costs of reproduction are mediated by male seminal fluid proteins (Sfps) transferred to females in the ejaculate (Wigby & Chapman 2005; Gioti *et al*. 2012; Perry *et al*. 2013). In particular, sex peptides (SP) are harmful proteins that alter female fecundity and decrease their sexual receptivity, with important carryover effects on their immunity, lifespan and fitness (Wigby & Chapman 2005; Gioti *et al*. 2012; Perry *et al*. 2013). Studies conducted in other insects show that temperature can influence male fertility via damage to the testes and/or sperm (Rohmer *et al*. 2004; Jørgensen *et al*. 2006; Lacoume *et al*. 2007; Nguyen *et al*. 2013). Similarly, two studies in other *Drosophilid* species found that temperature affected sperm motility and fertility, and ultimately caused partial sterility (Rohmer *et al*. 2004; Jørgensen *et al*. 2006). In this context, it seems reasonable to speculate that temperature might also impact negatively on other components of the ejaculate, for example on the amount or functionality of ejaculate components related with male harm to females (e.g. SP sex peptide; (Wigby & Chapman 2005)). This would explain why the generally high mating frequency at 29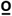C does not translate into significantly higher costs for females, but rather the opposite. It would not, however, explain why male harm levels at 21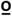C tended to be higher than at 29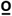C, despite lower net sexual harassment, re-mating rates, and lower increases of both these variables in a high sexual conflict scenario (Fig. 2–4).

To conclude, what transpires from our behavioural results is that no single mechanism seems to adequately explain why male harm increases more at 25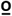C than at lower or higher temperatures. This suggests that harm to females via different mechanisms (e.g. sexual harassment and mating costs) is optimal at 25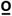C, and that deviations from this temperature affect underlying proximate mechanisms differently depending on the direction of this deviation. We also found that temperature modulates male harm effects on reproductive senescence and general reproduction costs (i.e. on actuarial senescence; Fig. 1) in a non-linear way, suggesting complex interactions with underlying proximate mechanisms. Disentangling these complex interactions certainly promises to be a challenging but exciting novel avenue of research that could reap valuable information on the dynamic interplay between temperature, ejaculate production and composition, and ageing processes.

## Acknowledgements

PC was supported by a “Ramón y Cajal” research fellowship (RYC-2013–12998). RGR and the research described here were supported by a MINECO Excelencia (CGL2017–89052-P to PC) co-funded by the European Regional Development Fund (FEDER), a SEJI research grant by the Generalitat Valenciana (SEJI/2018/037 to PC) and a 2018 Leonardo Grant for Researchers and Cultural Creators from the BBVA Fundation (to PC). VC was supported by a Ministerio de Educación Cultura y Deportes undergraduate collaboration grant (Beca de Colaboración 2017/2018).

